# Preserved suppressive function despite loss of Foxp3: insights into the identity of regulatory T cells

**DOI:** 10.1101/2025.01.17.633544

**Authors:** B Luckerbauer, AV Villarino, A Laurence, K Hirahara, G Sciume, Y Mikami, HY Shih, HW Sun, A Tosevska, M Brinkmann, SR Brooks, J Scheffel, F Meylan, B Afzali, M Bonelli

## Abstract

Regulatory T (Treg) cells are essential for maintaining immune homeostasis, with Foxp3 acting as the master transcription factor governing their differentiation and function. The acquisition of effector signatures in Treg cells is closely tied to the surrounding tissue-specific immune environment and typically occurs alongside Foxp3 expression. In this study, we investigated the transcriptomic and functional consequences of Treg-mediated regulation in a Th2-driven disease setting. The application of both *in vitro* systems and *in vivo* disease models allowed us to mimic Th2-mediated environments.

We could demonstrate Th2-driven loss of Foxp3 expression in Treg cells *in vitro* and *in vivo*. Transcriptomic analysis revealed a maintained Treg signature despite the loss of active Foxp3 expression. Functional characterization of Tregs both *in vitro* and *in vivo* uncovered a preserved suppressive capacity even in the absence of Foxp3.

Our findings unveil that, despite loss of Foxp3, a preserved Treg signature remains intact enabling the regulation of Th2-mediated diseases. The persistence of this regulatory transcriptome highlights the importance for developing Treg-cell therapy strategies in cancer and autoimmune diseases independent of Foxp3 expression.

## INTRODUCTION

Numerous lines of evidence highlight the forkhead family protein, Foxp3, as the master transcription factor (TF) governing various types of CD4^+^ regulatory T cells (Tregs) (1, 2) including thymic-derived Tregs (tTregs), peripherally-derived Tregs (pTregs) and *in vitro*-induced (iTreg) cells (3). Foxp3 executes its functional programs by binding to over 1000 genes, acting as a transcriptional activator or repressor (4). Treg populations can be differentiated on the basis of the methylation status of the CpG site-enriched CNS2 enhancer, containing the binding site for Foxp3 (5). While in tTreg cells, the CpG sites are fully demethylated, ensuring sustained Foxp3 expression, iTreg cells show diminished demethylation. Thus, causing an unstable expression of Foxp3 (6). Nevertheless, loss of Foxp3 expression in tTreg populations may still occur in response to inflammatory stimuli (7). This has led to a shift in classification of Treg cells with molecular characterization of Treg subsets revealing populations defined by co-expression of transcription factors specific to Th cell lineages (8, 9). Upregulation of selective lineage-determining TF associated with effector T cells generate functionally distinct subsets. Treg specific deletion of these master regulators such as Gata3 or T-bet unexpectantly results in spontaneous inflammatory disorders in mice (10, 11). Thus, it is essential that Treg cells gain a transcriptional effector program that allows to express certain master regulators (Gata3, T-bet, Rorγt) while repressing the rest of effector signatures to prevent spontaneous inflammation (12, 13). The intricate balance in the Treg transcriptional effector program highlights the importance of both intrinsic mechanisms and extrinsic signals in the fine-tuning of regulatory responses (14). This interplay is exemplified by the unique functionality of lineage-committed, microbiota-dependent pTreg cells, which demonstrate their immune regulatory capacities independently of Foxp3 (15). The functional independence of pTreg cells from Foxp3 underscores a critical distinction between the traditional view of Foxp3 as a master regulator and the broader network of mechanisms governing Treg cell differentiation and function. Emerging evidence supports this notion as a TF network consisting of Eos, Irf4, Satb1, Lef1 and Gata1 can not only enhance the binding of Foxp3 to its genomic targets, but can similarly activate the majority of the Treg signature, independently of Foxp3 (16), suggesting a genetic lock in the transcriptome of Treg cells (17, 18). Thus, Treg cells may retain their anti-inflammatory function, even after the loss of Foxp3 expression.

In this work we show that both tTreg and iTreg cells lose Foxp3 expression in response to antigen stimulation in the presence of IL-4, yet retain their ability to suppress activated effector T cells both *in vitro* and *in vivo*. Transcriptomic signatures of Treg and ex-Treg cells show a maintained phenotype despite the loss of Foxp3 expression.

## METHODS

### Mice and T cell culture

C57BL/6J, *Foxp3eGFP* and *Rag2*^*-/-*^ mice were purchased from Jackson Laboratory. *C57BL/6J Foxp3-GFPCre-Rosa26RFP* mice were kindly provided from Jeffrey A. Bluestone. All animal studies were performed according to the NIH guidelines for the use and care of live animals and were approved by the Institutional Animal Care and Use Committee of NIAMS. All cells were cultured in RPMI medium with 10% (vol/vol) FCS, 2 mM glutamine, 100 IU ml^-1^ of penicillin, 0.1 mg ml^-1^ of streptomycin and 20mM HEPES buffer, pH 7.2-7.5 (all from Invitrogen) and 2 mM β–mercaptoethanol (Sigma-Aldrich).

### Cell isolation and differentiation

CD4^+^ T cells from spleens and lymph nodes of 6–8-week-old mice were purified by negative selection and magnetic separation (Miltenyi Biotec) followed by sorting of naïve CD4^+^CD62L^+^CD44^-^CD25^-^GFP^-^ population using FACSAria III (BD).

For **iTreg** cell differentiation, naïve CD4 T cells were activated by plate-bound anti-CD3 (10 μg ml^-1^; eBioscience) and anti-CD28 (10 μg ml^-1^; eBioscience) in media for 3 days with human IL-2 (100 IU ml^-1^), human TGF-β1 (10 ng ml^-1^, R&D Systems), anti-IL-4 neutralizing antibodies (10 μg ml^-1^, BioXCell) and anti-IFN-γ neutralizing antibodies (10 μg ml^-1^, BioXCell). After 3 days, Foxp3^+^GFP^+^ iTreg cells were sorted, activated by plate-bound anti-CD3 (1 μg ml^-1^) and anti-CD28 (1 μg ml^-1^). After 3 days, Foxp3^+^GFP^+^ iTreg cells were resorted to ensure purity.

For **ex-iTreg** cell differentiation, iTreg cells were generated first as described above for 3 days. After 3 days, sorted Foxp3^+^GFP^+^ iTreg cells were activated by plate-bound anti-CD3 (1 μg ml^-1^) and anti-CD28 (1 μg ml^-1^) in media with IL-4 (20 ng ml^-1^). After 3 days, Foxp3^-^GFP^-^ ex-iTreg cells were resorted (ori. iTreg).

For **Th2** cell polarization, naïve CD4 T cells were activated by plate-bound anti-CD3 (10 μg ml^-1^) and anti-CD28 (10 μg ml^-1^) in media for 3 days with human IL-2 (100 IU ml^-1^), IL-4 (20 ng ml^-1^, R&D Systems) and anti-IFN-γ neutralizing antibodies (10 μg ml^-1^). After 3 days, Foxp3^-^GFP^-^ Th2 cells were sorted and activated by plate-bound anti-CD3 (1 μg ml^-1^) and anti-CD28 (1 μg ml^-1^) in media with IL-4 (20 ng ml^-1^). After 3 days, Foxp3^-^GFP^-^ Th2 cells were re-sorted.

### Flow cytometry

The following antibodies were used for surface and intracellular staining. For cell surface staining: anti-CD4 (clone: RM4-5), anti-CD25 (clone: 7D4), anti-CD45.1 (clone: A20), anti-IL-2 (clone: JES6-5H4), anti-IL-10 (clone: JES5-16E3) and anti-CTLA-4 (UC10-4F10-11) were purchased from BD Bioscience. Anti-CD44 (clone: IM7), anti-CD62L (clone: MEL-14), anti-Foxp3 (clone: FJK-16s), anti-CD45.2 (clone: 104), anti- and anti-GITR (clone: DTA-1) were purchased from eBioscience. were purchased from Biolegend. Expression of TFs was assessed by intracellular staining. For intracellular cytokine staining, cells were stimulated for 4 hours with Phorbol 12-myristate 13-acetate (PMA 50 ng ml^-1^) and ionomycin (500 ng ml^-1^) with the addition of brefeldin A (GolgiPlug; BD). Afterwards, cells were fixed with 4% formyl saline, permeabilized with 0.1% saponin buffer and stained with fluorescent antibodies before analyzing on a FACS Verse (BD). Events were collected and analyzed with Flow Jo software (Tree Star).

### RNA Sequencing

RNA Sequencing was performed as described previously (32). Total RNA was prepared from approximately 1 million cells by using mirVana miRNA Isolation Kit (AM1560, ABI). 200 ng of total RNA was subsequently used to prepare RNA-seq library by using TruSeq SR RNA sample prep kit (FC-122-1001, Illumina) by following manufacturer’s protocol. For the ex vivo analysis, two biological replicates per condition (Foxp3+, exFoxp3, nonFoxp3) were tested. The libraries were sequenced for 50 cycles (single read) with a HiSeq 2000 (Illumina). Raw sequencing data were pseudo-aligned the bulk mRNA-seq reads to the mouse transcriptome (ENSEMBL release 96, GRCm38 cDNA) using kallisto (v0.48) (33) to estimate transcript abundance. Transcripts were then collapsed to obtain the gene level counts per sample. Data was normalized and differentially expressed genes were identified using DESeq2 (q-value < 0.05; absolute fold change >= 2; mean count higher than 10) (34). *In vitro* RNA-seq samples were mapped to the GRCm39 using We also aligned nascent RNA-seq reads to the mouse genome (STAR v2.7.8a) (35), and data was further normalized and processed using DESeq2 as described. Principal component analysis was performed using PCAtools v2.16.0 [10.18129/B9.bioc.PCAtools]. Gene signatures were defined as gene significantly upregulated compared to non-Foxp3+ cells, using the aforementioned criteria. The nVennR v0.2.3 package was used to define overlapping gene signatures.

### Enrichment analysis

Differentially expressed genes as defined previously were then functionally enriched using the enrichR package in R and clustered and summarized using SummArIzeR v0.01 [http://10.5281/zenodo.14627791]. Briefly, the databases used for enrichment analyses were as follows: “GO_Biological_Process_2023”,”WikiPathways_2024_Mouse”, “MSigDB_Hallmark_2020”, “Mouse_Gene_Atlas” from enrichR. Top 3 terms per database and comparison were selected and terms with fewer than 5 genes were filtered out. Genes were split by up/downregulation in figures 1g and 2e. The term similarity scores were then calculated based on Jaccard similarity of genes and clusters were inferred from the similarity matrix using a Walktrap community finding algorithm, Edges with a threshold below 0.32 were removed from further analysis in figures 1g and 2e. A threshold of 0.15 was used for (Figure 4a).

**Figure 1.**
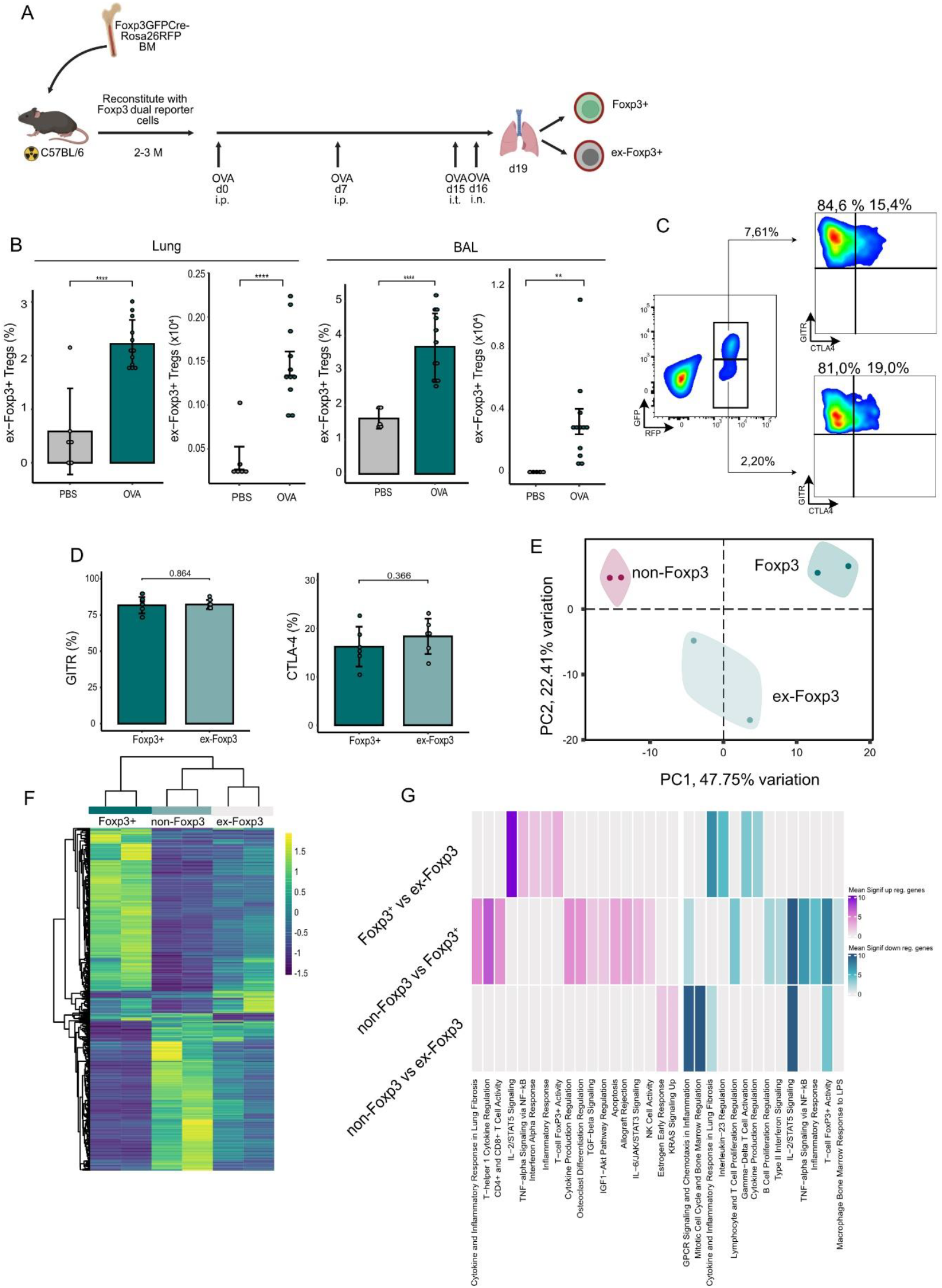
Maintained Treg phenotype despite loss of Foxp3 expression at inflammatory site. **a**) Schematic representation of the study design and interventional timepoints. Irradiated C57BL/6 mice were reconstituted with Foxp3-GFPCre-Rosa26RFP reporter cells and subjected to ovalbumin (OVA)-induced asthma. **b)** T cells were isolated from the lung and bronchoalveolar lavage (BAL) fluid from asthma induced and control mice and analyzed for the expression of RFP+GFP+ and RFP+GFP-cells. Bar charts depict pooled data from 2 independent experiments (mean value ± SEM). **c)** T cells were isolated from the lung of OVA and PBS treated mice and both CD4^+^RFP^+^GFP^+^ and CD4^+^RFP^+^GFP^-^ cells were stained for the surface expression of GITR and CTLA-4. Representative plots are shown. **d)** Bar charts indicate percentage of GITR and CTLA-4 expression in Foxp3+ and ex-Foxp3 cells (mean value ± SEM) **e)** Principal component analysis (PCA) of the global expression measured by RNA-seq. Samples are labeled by cell type. **f)** Heatmap of the the differentially expressed genes between all three cell types using a pairwise analysis. Values depict scaled normalized read counts. Differentially expressed genes were defined as those with an FDR adjuster p-value (q-value) below 0.05 and an absolute fold change of 2. **g)** Functional enrichment analysis for pairwise differentially expressed genes between the three cell populations, separated byup- and down-regulated genes. The heatmap depicts pooled adj. p values for clustered terms derived from the “GO_Biological_Process_2023”,”WikiPathways_2024_Mouse”, “MSigDB_Hallmark_2020”, “Mouse_Gene_Atlas” databases. Means were compared using a Student’s t-test. Significance thresholds were defined as follows: *P < 0.05, **P < 0.01, ***P < 0.001, ****P < 0.0001.

### *In vitro* suppression assay

FACS purified Foxp3GFP^+^ iTreg, Foxp3GFP^-^ ex-iTreg cells, were cultured for 3 days without or with IL-4, respectively. Naïve CD4+ T cell were subjected to Th2 polarizing conditions to isolate Foxp3GFP^-^Th2 cells. A co-culture of responder T cells (5 × 10^4^ CFSE-labeled naïve CD45.1^+^CD4^+^CD62L^+^ CD44^low^), antigen-presenting cells (1×10^4^ CD11c^+^ dendritic cells isolated by immunomagnetic selection (Miltenyi)) and sort purified Foxp3GFP^+^ iTreg, Foxp3GFP^-^ ex-iTreg and Foxp3GFP^-^Th2 cells was conducted in a 96-well round-bottom plate. A serial dilution of Foxp3GFP^+^ iTreg, Foxp3GFP^-^ ex-iTreg and Foxp3GFP^-^Th2 cells from the ratios 1:1 to 1:16 ensured investigations of suppressive behavior under varying Treg:responder cell concentrations. Cells were stimulated with soluble 1 μg ml^−1^ anti-CD3 antibody for 3 days at 37 °C and 5% CO_2_. Proliferation of responder cells was measured by flow cytometry.

### Adoptive transfer colitis

*In vivo* colitis model was done as described previously (36). Briefly, *Rag2*^*-/-*^ mice were injected intravenously with 4 × 10^5^ FACS-sorted wild-type naive CD45.1^+^CD4^+^CD25^-^ CD45Rb^hi^ cells with or without 2 × 10^5^ iTreg, ex-iTreg or Th2 cells. Mice were monitored weekly for weight loss and signs of disease. 8 weeks post adoptive transfer, samples for histology sections were taken from the colon descendens, fixed in formaldehyde and stained for hematoxylin and eosin.

### Ova-Induced lung inflammation

In order to fate-map Foxp3, bone marrow cells from *Foxp3-GFPCre-Rosa26RFP* were harvested, whereof 1 × 10^7^ cells were injected in wild type C57BL/6 mice conditioned with total-body irradiation of 950 cGy. For induction of asthma, CD45.1^+^ or Foxp3 dual reporter reconstituted mice were sensitized systemically via a 200-μl intraperitoneal injection containing either 100 μg chicken OVA (Sigma, St Louis, MO) or PBS emulsified in an equal volume mixture with alum (Thermo Scientific, Waltham, MA) on days 0 and 7. On day 13, 1 × 10^6^ *in vitro* differentiated iTreg, ex-iTreg or Th2 cells were injected in *CD45*.*1*^*+*^ mice. All mice were challenged with 100 μg OVA or PBS per 30 μl inoculum intratracheally on day 15 and intranasally on day 16. Mice were euthanized on day 19. Lungs were perfused with PBS and lung cell preps were obtained by incubating lung fragments with 100 U collagenase for 1 h. Cells were analyzed for the expression of CD4, CD45.1, CTLA-4, GITR and Foxp3.

### Statistics

For statistical analysis, Student’s *t*-test, Anova tests and Hypergeometric tests were used unless otherwise specified.

## RESULTS

### Maintained Treg phenotype despite loss of Foxp3 expression at inflammatory site

Since previous work has shown that Treg cells can lose Foxp3 expression and acquire effector cell function (19-22) under inflammatory conditions, we performed an Ovalbumin (OVA)-induced asthma model in a dual color reporter mouse strain in which prior expression of Foxp3 can be fate mapped *in vivo* (23, 24) analyzed Treg cells from lungs **(Figure 1a)**.

At the site of inflammation, we observed a statistically significant increase in RFP^+^GFP^-^ ex-Foxp3 cells in both the inflamed lung tissue and bronchoalveolar lavage (BAL) fluid in the OVA experimental group compared to PBS **(Figure 1b)**. In the identified ex-Foxp3 cell population, expression of Treg associated markers such as CTLA-4 and GITR were maintained **(Figure 1c-d)**, suggesting that Treg associated gene expression can be maintained.

To assess the transcriptional patterns underlying these changes, we performed RNA-seq on Foxp3+, ex-Foxp3 and non-Foxp3 sorted cell populations. Principal component analysis revealed distinct clusters corresponding to the three populations, highlighting the transcriptional difference between Foxp3+ and ex-Foxp3 cell populations **(Figure 1e)**. To better understand the relationship among the 3 cell populations, unsupervised hierarchical clustering was performed on the pre-defined differentially expressed genes (adjusted p-value < 0.05, absolute fold change > 2). The analysis indicated ex-FoxP3 cells represent a hybrid state with features of both Foxp3+ and non-Foxp3 cells **(Figure 1f)**. Differential expression between Foxp3+ and ex-Foxp3 cell populations revealed an upregulation of genes associated with IL-2/STAT5 Signaling and T-cell Foxp3 activity, while a downregulation in genes associated with inflammatory response was observed **(Figure 1g)**.

Together, these data indicate a transcriptomic similarity between Foxp3+ and ex-Foxp3 cell populations, which are generated in a Th2-mediation inflammation.

### Preservation of the Treg transcriptomic signature in ex-iTreg cells

Since the loss of Foxp3 in Treg cells occurred in the setting of lung inflammation, we established an *in vitro* system to examine the consequences of Th2 cytokine driven destabilization of Foxp3 expression in Treg cells **(Figure 2a)**. We sought to explore the transcriptional relationship between cell populations derived from the *in vitro* setting and found similar gene expression patterns as described in the Th2-driven inflammation model **(Figure 2b)**. Hierarchical clustering analysis positioned the transcriptomic expression profile of ex-iTreg cells as intermediate between classical iTreg cells and effector Th2 populations **(Figure 2c)**. Expression of Treg signature genes revealed similarities of ex-iTreg and Th2 cells, distinct to iTreg cells. While only iTreg cells could be identified by Foxp3 expression, ex-iTreg cells follow the gene expression pattern of Th2 cells **(Figure 2 d)**, in line with the *in vivo* findings. Differential gene expression analysis between iTreg and ex-iTreg cells revealed an upregulation in Foxp3 activity, in contrast to a downregulation of cytokines and inflammatory responses **(Figure 2e)**.

**Figure 2.**
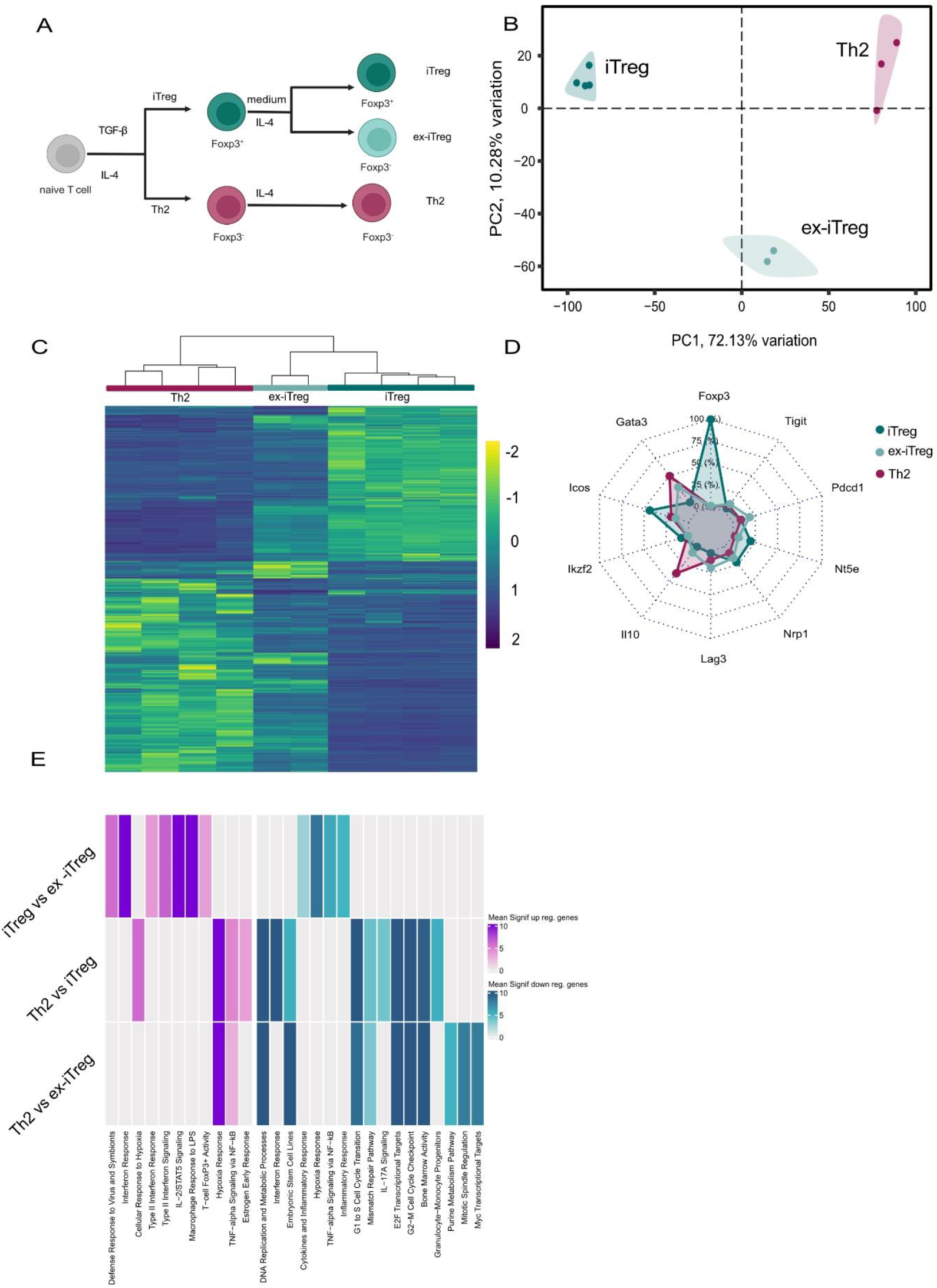
Preservation of the Treg transcriptomic signature in ex-iTreg cells. **a)** Naive CD4+ T cells were subjected to in vitro polarizing conditions towards iTreg and Th2 cells, supplementing the media with TGFb or IL-4, respectively. Sorted Foxp3+GFP+iTreg cells were stimulated with IL-4 to generate Foxp3-GFP-ex-iTreg cells. **b)** Principal component analysis (PCA) of the global expression measured by RNA-seq. Samples are labeled by cell type. **c)** Heatmap of the the differentially expressed genes between all three cell types using a pairwise analysis. Values depict scaled normalized read counts. Differentially expressed genes were defined as those with an FDR adjuster p-value (q-value) below 0.05 and an absolute fold change of 2. **d)** Combined radar chart comparing expression pattern of the top 10 Treg-associated genes across the iTreg, ex-iTreg and Th2 cell populations. Values were normalized (min-max normalization in percent) **e)** Functional enrichment analysis for pairwise differentially expressed genes between the three cell populations, separated by up- and down-regulated genes. The heatmap depicts pooled adj. p values for clustered terms derived from the “GO_Biological_Process_2023”,”WikiPathways_2024_Mouse”, “MSigDB_Hallmark_2020”, “Mouse_Gene_Atlas” databases.

Collectively, these data demonstrate that *in vitro* exposure of iTregs to IL-4 is sufficient to recapitulate the described *in vivo* phenomenon of Th2-mediated Foxp3 loss.

### Retention of suppressive functionality in ex-iTreg cells

In addition to the transcriptional signature Foxp3^+^ Treg cells are defined by their suppressive capacity, which can be tested in classical *in vitro* and *in vivo* suppression assays (25, 26). We therefore investigated whether the retention of a characteristic gene expression pattern was associated with suppressive function in the absence of Foxp3. Therefore, in an established *in vitro* system iTreg, IL4-induced ex-iTreg und Th2 cells were generated, allowing for the division of iTreg cells in two distinct populations based on Foxp3 expression by cell sorting **(Figure 3a)**. First, we compared all three cell populations in a classical *in vitro* suppression assay. These studies revealed that in contrast with Th2 cells, IL-4 induced ex-iTreg cells, were equally potent inhibitors of naïve T cell proliferation, compared with their Foxp3^+^ precursors **(Figure 3b-c)**. We next investigated if the maintenance of suppressive capacity of IL-4 induced ex-iTreg cells was evident in an *in vivo* adoptive T cell transfer model of colitis. For these studies, congenic naïve T cells were co-transferred with either iTreg, Th2 or IL-4 induced ex-iTreg cells into Rag2^-/-^ mice, which were monitored for disease progression. Consistent with our *in vitro* findings, we observed that unlike Th2 cells, both iTreg and IL-4 induced ex-iTreg cells were able to limit weight loss **(Figure 3d)**, intestinal pathology **(Figure 3e)** and severity of inflammation **(Figure 3f + g)**. Re-isolated intestinal Treg cells of the previously administered IL-4 induced ex-iTregs remained Foxp3-negative, even at 8 weeks post-transfer. **(Figure 3h)**.

**Figure 3.**
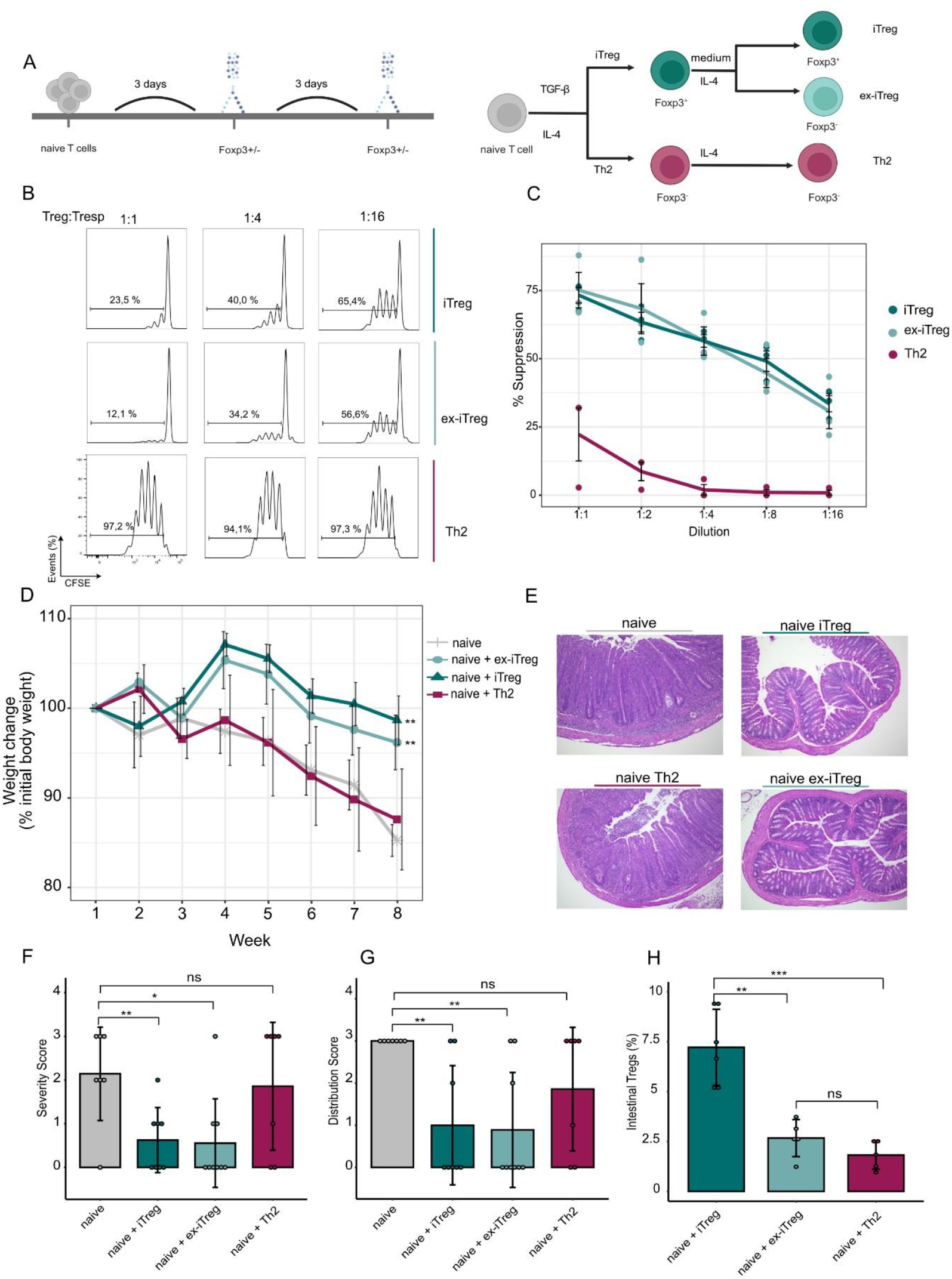
Retention of suppressive functionality in ex-iTreg cells. **a)** Naive CD4+ T cells were subjected to in vitro polarizing conditions towards iTreg and Th2 cells in a 3-day cell culture. Sorted Foxp3+GFP+iTreg cells were either stimulated with IL-4 to generate Foxp3-GFP-ex-iTreg cells or cultured in standard RPMI medium. **b)** In vitro suppression assays. CFSE labeled naïve cells were cultured without or with the indicated ratios of iTreg, ex-iTreg or Th2 cells. Cell proliferation was measured by CFSE dilution. A representative experiment is shown. **c)** Graph depicts 3 independent in vitro suppression assays; percentage of naïve T cell suppression is shown (mean ± SEM). **d)** In vivo adoptive T cell transfer colitis to evaluate iTreg, ex-iTreg and Th2 cells suppressive capacity. Presence of autoimmune disease was assessed by changes in body weight over time (% to initial body weight). **e)** Representative micrographs of colones descendentes are shown. **f-g)** Histological scores as severity and distribution of inflammation are depicted (n=4 independent experiments, mean ± SEM) based on data from figure 3e. **h)** On week 8, cells were re-isolated from colon descendens and analyzed for Foxp3 expression. Bar graphs show percentages of cells expressing Foxp3 assessed by flow cytometry (mean ± SEM). Means were compared using a Student’s t-test. Significance thresholds were defined as follows: *P < 0.05, **P < 0.01, ***P < 0.001, ****P < 0.0001.

These data indicate that, despite the loss of Foxp3 expression, IL-4 induced ex-iTreg+ cells preserved the ability to exert suppressive function *in vitro* and *in vivo*.

### Th2-mediated inflammation destabilizes Foxp3 expression in iTregs

As loss of Foxp3 was not accompanied by diminished suppressive capacity, we investigated the similarities between ex-Foxp3 and Foxp3+ Tregs *in vivo* and *in vitro*. An overlap of gene signatures was identified between Foxp3+ and ex-Foxp3, as well as iTreg and ex-iTreg cells **(Figure 4a)**. Hierarchical clustering connected the gene expression of lung-derived Foxp3+ Treg cells to the generated iTreg cells based on similar signature genes associated with IL2/STAT5 signaling as well as inflammatory response. The gene expression signature derived from the overlap between iTregs and ex-iTregs could be linked to the unique signature of lung-derived ex-Foxp3 Tregs, clearly separating this signature from their Foxp3+ counterparts, which showed a higher enrichment of classical Treg expression profile **(Figure 4b)**.

**Figure 4.**
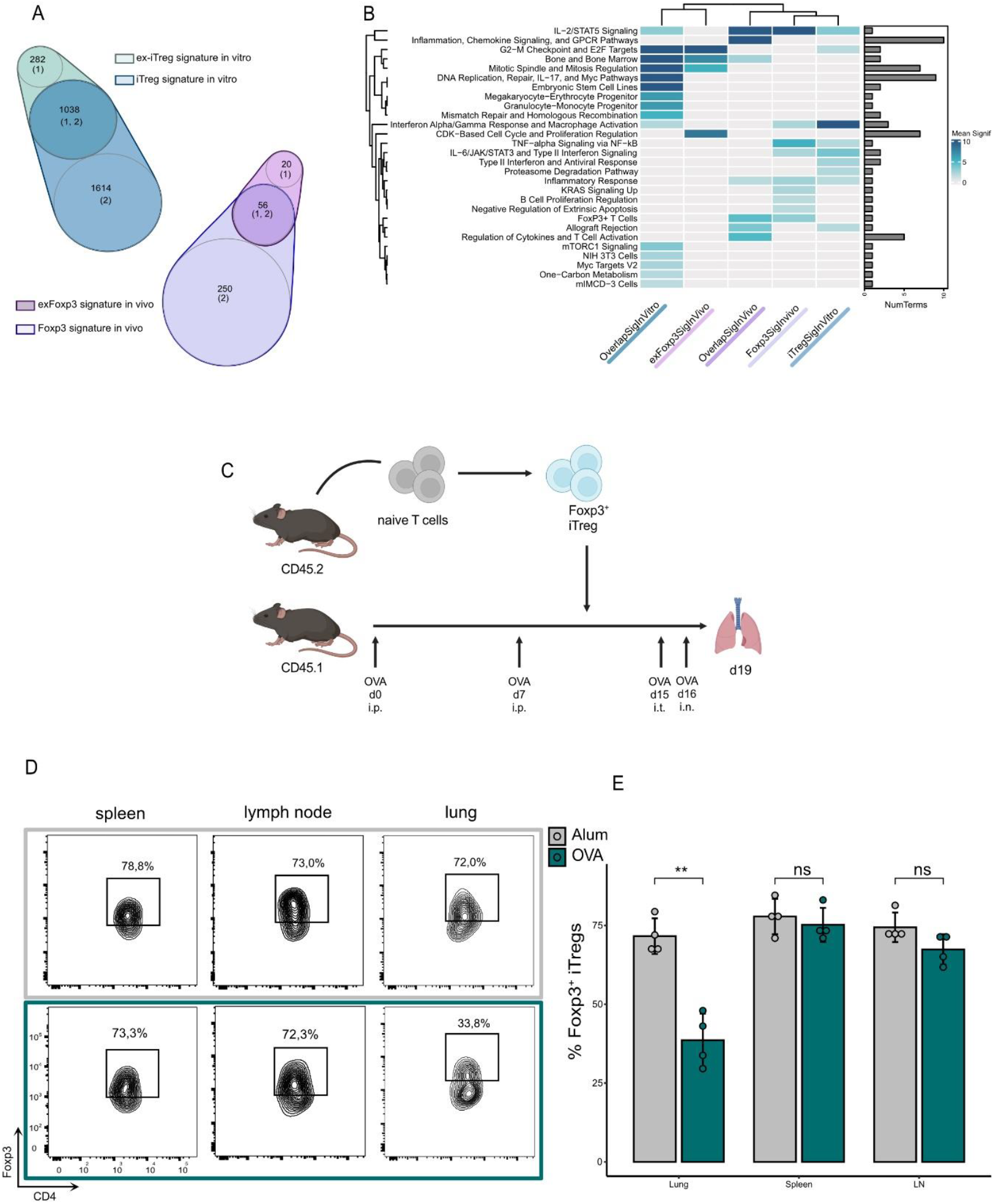
Th2-mediated inflammation destabilizes Foxp3 expression in iTregs. **a-b)** Functional enrichment analysis for unique and shared signature genes between Foxp3 and exFoxp3 cells, upregulated compared to non-Foxp3 both in vivo and in vitro. The number of unique and overlapping genes for each signature are depicted in the Venn Diagrams. The heatmap depicts pooled adj. p values for clustered terms derived from the “GO_Biological_Process_2023”,”WikiPathways_2024_Mouse”, “MSigDB_Hallmark_2020”, “Mouse_Gene_Atlas” databases. **c)** Schematic representation of the study design and interventional timepoints. Ovalbumin (OVA)-induced asthma model was performed using mice exhibiting a CD45.1 background. Simultaneous culture of iTregs exhibiting a CD45.2 background allowed for administration of iTregs into the OVA-induced asthma model before intra-tracheal intervention. **d)** Foxp3 expression of administered CD45.2+ iTreg cells harvested from spleen, lymph nodes and lung of OVA-treated and control mice was measured by flow cytometry. A representative experiment is shown. d) Bar graphs indicate percentage of CD45.2+ iTregs actively expressing Foxp3 in spleen, lymph nodes and lung in both OVA-treated and control mice, as quantified from figure **4c**. Means were compared using a Student’s t-test. Significance thresholds were defined as follows: *P < 0.05, **P < 0.01, ***P < 0.001, ****P < 0.0001.

As iTregs cells exhibited a classical Treg transcriptomic profile *in vitro*, we next sought to investigate their fate and function in the context of a Th2-associated inflammatory disease *in vivo*. Therefore, we immunized and challenged CD45.1^+^ mice with OVA. The iTreg cells were transferred after immunization, re-isolated after challenge and analyzed for the expression of Foxp3 by flow cytometry **(Figure 4c)**. We found pronounced loss of Foxp3 expression following adoptive transfer of iTreg cells, but only at the site of inflammation (lung) **(Figure 4d-e)**.

Our data demonstrates that Treg cells adapting to a Th2-mediated inflammatory environment can sustain regulatory function as they transition to an effector-like phenotype despite downregulation of Foxp3.

## DISCUSSION

In this study, we demonstrated that Th2-driven inflammatory conditions can result in the loss of Foxp3 expression in Treg cells, that is not affecting their suppressive function. This population of Treg cells maintained a characteristic regulatory transcriptomic signature, without actively expressing Foxp3, both *in vitro* and *in vivo*.

The acquisition of effector signatures in Treg cells, closely aligned with the surrounding tissue-specific effector environment, is a well-documented phenomenon that occurs concurrently with Foxp3 expression (8, 27-29). Despite growing evidence that other transcription factors contribute to shaping the transcriptomic profile of Treg cells, Foxp3 remains recognized as the master regulator of their identity and function (15, 16).

Our findings challenge this paradigm by revealing that in an OVA-induced asthma model, Treg cells lose active Foxp3 expression within sites of established Th2-driven inflammation. Remarkably, despite loss of their canonical master regulator, this cell population retains its regulatory phenotype, as evidenced by the persistence of a regulatory transcriptome both *in vitro* and *in vivo*.

This observation raises intriguing questions about the preserved transcriptomic signature underpinning Treg function and stability. The maintenance of a regulatory transcriptome in the absence of active Foxp3 expression suggests that alternative transcriptional programs may stabilize the regulatory identity of these cells. Prior studies have indicated that Foxp3-independent regulatory pathways, including from TFs such as Blimp-1 and Helios, can sustain Treg functionality under certain conditions (30, 31). Our data lend further support to the notion that Treg cells possess a degree of plasticity, which, even despite the loss of Foxp3, allows them to adapt to highly inflammatory environments while preserving their suppressive capacity.

Despite the novel insights provided by our findings, a limitation to be considered is the poorly understood mechanism of Foxp3 downregulation within Th2-dominant inflammatory environments and whether this phenomenon extends to other inflammatory conditions.

This presence of a regulatory transcriptome in cells that have lost their cognate master regulator has important implications for manipulating Treg cells in cancer and autoimmunity and may serve as an indicator of cellular identity independently of Foxp3 expression. Thus, therapies based on the adoptive transfer of Treg cells may benefit from employing measuring strategies that offer more insight into the regulatory potential at the site of inflammation than stable Foxp3 expression.

## Data availability statement

Data will be made available on acceptance.

## Competing financial interests

The authors declare no competing financial interests.

## Acknowledgements

We thank J. Simone, J. Lay and K. Tinsley (Flow Cytometry Section, NIAMS), G. Gutierrez-Cruz (Office and Science and Technology, NIAMS), the NIAMS LACU staff for their excellent technical support and A. Poholek, G. Vahedi, K. Singleton, K. Jiang and O. Engel for supporting experiments and analysis. We thank Victoria Hoffmann for expert help in histology scoring. This work was supported by the Intramural Research Programs of NIAMS. We thank JJ O’Shea, J Bluestone and Y Kanno for providing conceptual and intellectual input.

## Author contributions

M.B., A.L., A.V., MaB, AT and BL designed experiments, interpreted data and wrote the manuscript. M.B., A.V., K.H., G.S, H-W.S., H.S. J.S., Y.M., A.B, E.H., T.E, F.M. and Y.K. performed experiments. M.B., AT, MaB, BL, S.P., S.B., H-W.S. and H.S. analyzed genomic data. F.C. made helpful suggestions.

## Notes

### Competing Interest Statement

The authors have declared no competing interest.

